# The rate of transient beta frequency events predicts impaired function across tasks and species

**DOI:** 10.1101/128769

**Authors:** Hyeyoung Shin, Robert Law, Shawn Tsutsui, Christopher I. Moore, Stephanie R. Jones

**Affiliations:** Department of Neuroscience, Brown University, Providence, Rhode Island; Center for Neurorestoration and Neurotechnology, Providence VA Medical Center, Providence, Rhode Island

## Abstract

Beta frequency oscillations (15-29Hz) are among the most prominent signatures of brain activity. Beta power is predictive of many healthy and abnormal behaviors, including perception, attention and motor action. Recent evidence shows that in non-averaged signals, beta can emerge as transient high-power “events”. As such, functionally relevant differences in averaged power across time and trials can reflect accumulated changes in the number, power, duration, and/or frequency span of the events. We show for the first time that functionally relevant differences in averaged prestimulus beta power in human sensory neocortex reflects a difference in the number of high-power beta events per trial, i.e., the rate of events. Further, high power beta events close to the time of the stimulus were more likely to impair perception. This result is consistent across detection and attention tasks in human magnetoencephalography (MEG) and is conserved in local field potential (LFP) recordings of mice performing a detection task. Our findings suggest transient brain rhythms are best viewed as a “rate metric” in their impact on function, and provides a new framework for understanding and manipulating functionally relevant rhythmic events.

## Introduction

Beta frequency oscillations are among the most prominent signatures of brain activity. Modulations in beta power correlate with perceptual and motor demands ((1-10)) and abnormalities are used as a biomarker of neuropathology (e.g. Parkinson’s disease, Alzheimer’s disease, Autism Spectrum Disorder (11-13)). Yet, how and why beta impacts function is yet unknown. The temporal signatures of rhythmic activity may prove crucial to understanding their importance in brain function (14-16). The vast majority of studies reporting functional correlates of brain rhythms rely on smoothing the data within fixed frequency bands and averaging across time and/or trials, where the temporal dynamics of the rhythmic activity is lost.

Several recent studies have appreciated the fact that rhythmic activity is often transient in non-averaged data, lasting a few cycles (14, 17-21). Beta frequency activity (15-29Hz) has been shown to emerge as brief events typically lasting <150ms. Transient beta has been reported in many brain areas including somatosensory (17, 18, 22), motor (19), occipital (23), and frontal cortex (18, 20), and basal ganglia structures (19, 21, 24), and across recording modalities and species (18). Further, differences in the accumulated density of transient beta bursts across trials predict functionally relevant differences in averaged beta power reflecting motor and cognitive demands in monkey LFP data (19, 20).

We have previously shown that shifts in spatial attention modulate post-cue / prestimulus beta power, such that average power decreases in the attended SI location (10, 25). Correspondingly, lower prestimulus beta power (averaged in the 1-second prestimulus time window) predicts an increase in the probability of perception of tactile stimuli at perceptual threshold (1). These prior studies did not investigate the relationship between the transient nature of the beta events on specific trials and function, a highly relevant question given the episodic nature of beta, including our recent findings showing that prestimulus beta activity is transient (<150ms) in human MEG signals from primary somatosensory (SI) and frontal cortex (17, 18).

Here, we found that functionally relevant differences in averaged prestimulus beta power in human neocortex emerges from a difference in the number of high power beta events per trial (e.g. the rate of beta events). This result is conserved in both detection and attention tasks (1, 26): The rate of beta events in the attended location decreases, and a lower rate of prestimulus beta events predicts enhanced detection of threshold stimuli. Moreover, non-detected trials were more likely to have a high-power beta event within 300ms of the stimulus. These results are the first to show that differences in the rate and timing of prestimulus beta events underlies the correlation between beta power and behavior in humans. Results were found to be highly conserved in LFP recordings from awake behaving mice performing a similar tactile detection task. The strong parallel in the character of beta events across behavior, recording scales, and species indicates a fundamental consistency in the nature of this much studied dynamic in the mammalian neocortex.

## Results

### Mean prestimulus beta power is higher on non-detected and attend-out trials

We have previously shown that prestimulus beta rhythms in human SI are predictive of tactile perception, such that there is a negative relationship between beta power averaged 1 second prestimulus and the probability of detecting a brief tap to the contralateral finger (1). Figure 1A confirms this result, showing that rate of detection (i.e., hit rate), measured as percent change from mean (PCM), decreases as averaged 1 second prestimulus beta power increases. Here, we show that this result also holds for LFP recordings from the SI “barrel” neocortex of mice performing a similar vibrissa deflection detection task (Figure 1B). Further, extending our prior investigation of changes in SI beta power during a somatosensory cued-attention task (1, 10), we find a similar negative relationship between prestimulus / post-cue beta power and the percent of attend-in trials (Figure 1C). In agreement with these analyses, grand averaged prestimulus power for each subject was significantly different across behavioral outcomes and conditions (Figure 1ii; p<0.05, two sampled left-tailed Wilcoxon signed-rank test).

**Figure 1.**
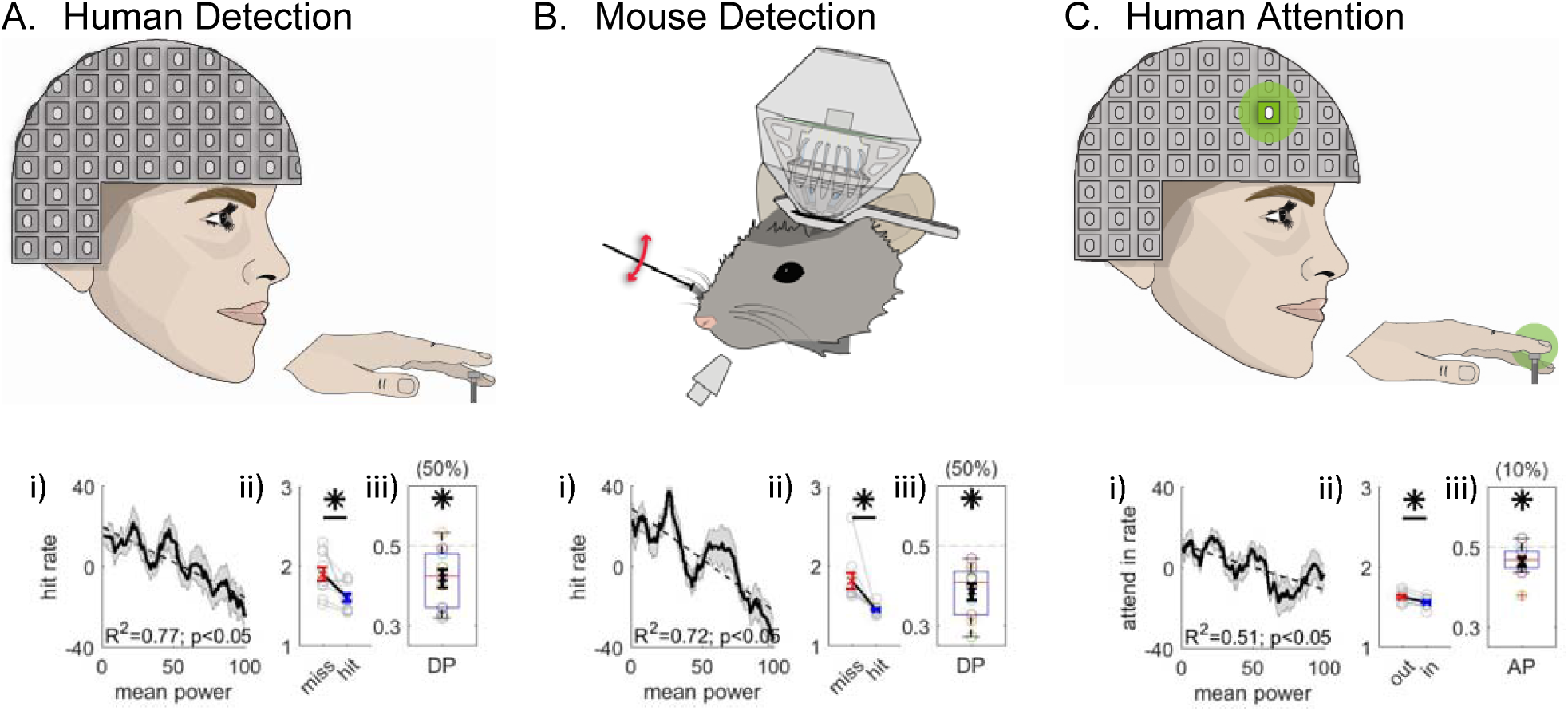
Average prestimulus beta (15-29Hz) power is higher on non-detected / attend-out trials. **i)** Mean (black), ±standard error of the mean (SEM, grey), and linear regression (dotted line) of averaged 1-second prestimulus SI beta band (15-29Hz) shows power negatively correlates with performance in **A.** human MEG recordings during a tactile detection task, **B.** mouse LFP during a vibrissae deflection detection task, and **C.** human MEG during a cued spatial attention task. Linear regression was performed for 21-trial bin percentile against detection rate / attend-in rate, and the *R*^2^ value and F-test p-values are shown. **ii)** Average beta power pooled across trials under each condition are shown for each subject / session (grey open circles), with the mean across subjects / sessions depicted in red and blue x marks and ±SEM as error bars. Averaged beta power was significantly larger in non-detected (miss) / attend-out trials (left-tailed Wilcoxon signed-rank test, asterisk p<0.05). **iii)** Box and whisker plots depict distribution of detect probability (DP) / attend probability (AP) across subjects / sessions. Mean and ±SEM across subjects are plotted with thick black x sign and error bars, respectively. One-sample left-tailed Wilcoxon signed rank test was applied to test whether the median across subjects / sessions was significantly less than 0.5. The numbers in parentheses at the top part of the plots shows the percentage of subjects / sessions with DP / AP significantly less than 0.5.

To further investigate the potential impact of prestimulus beta power on behavior, we conducted an ideal observer analysis for discriminating detected and attend-in trials versus non-detected and attend-out trials (27), respectively. These discrimination values are termed detect probability (DP) and attend probability (AP). A DP or AP value of 0.5 indicates no predictability, and <0.5 indicates that higher trial mean prestimulus beta power was predictive of non-detected or attend-out trials. Box plot distributions of these values across subjects or sessions are plotted in Figure 1iii. In each data set, the median of DP and AP distributions were significantly <0.5, such that higher power indicated the subject was less likely to detect or attend to the stimulus (Figure 1iii asterisks p<0.05, one-sample left-tailed Wilcoxon signed rank test; humans N=10, 1 session each, mice N=2, 5 sessions each). While this effect was significant when applying an across subject comparison, in an individualized bootstrap analysis the effect was stronger in the detection data sets, where in 50% of individual subjects and sessions high beta was predictive of non-detection, as compared to the attention data set where only 10% of the subjects showed a significant effect (Figure1iii, see parentheses above each plot, bootstrapping 10,000 times to determine 95% confidence interval. DP is significantly less than 0.5 if the upper boundary of the bootstrapped confidence interval is less than 0.5).

### Beta emerges as brief events on individual trials

The results in Figure 1 establish that prestimulus beta power is predictive of perception and attentional allocation. Typically, power is averaged across time, frequency and trials to investigate the relationship between brain rhythms and function. A key step in understanding how beta impacts function is to identify features in the unaveraged signal that contribute to this relationship. As discussed above, beta emerges as a transient surge in power on individual trials (i.e., events) (Figure 2ii). Because values in the spectral domain are non-negative, the accumulation of events across trials creates a continuous band of activity in the average, often misinterpreted as a sustained rhythm (Figure 2i; (14)). We confirm this observation for prestimulus LFP beta activity in mice during the vibrissa deflection detection task, and in the prestimulus and post-cue period during our cued attention task in humans (Figure 2B/C).

**Figure 2.**
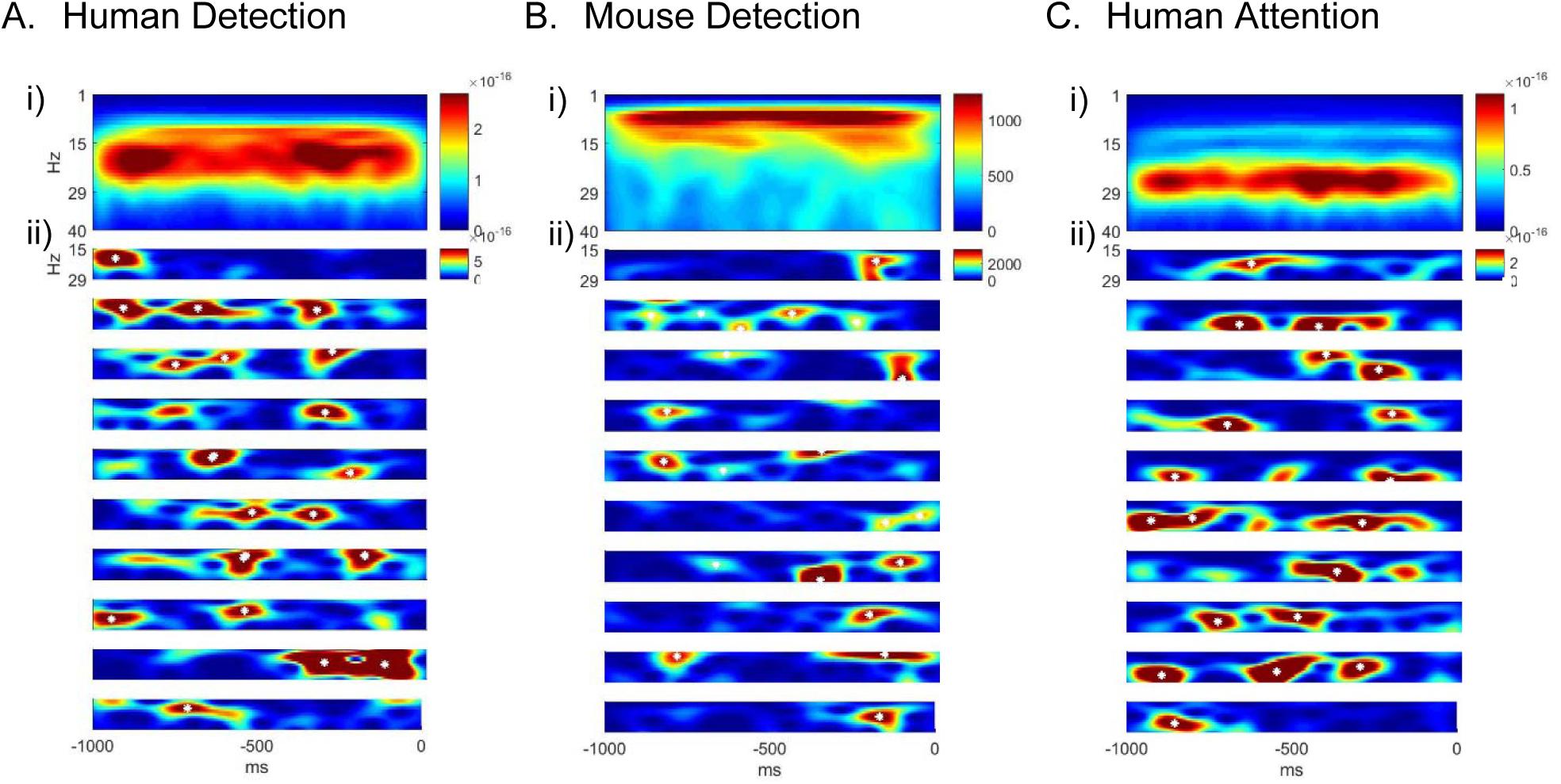
Beta emerges as brief events in non-averaged spectrograms. A. Top panel shows averaged spectrogram (1-40Hz) in the 1 second prestimulus period from 100 trials, from an example subject in the human detection dataset (average across 100 non-detected trials). The bottom panels show examples of prestimulus beta band (15-29Hz) activity in 10 trials from the same data set. White asterisks denote local maximas in the spectrogram with maxima power above 6X median power of the maxima frequency. **B.** Same format as A in mouse detection task (average across 130 non-detected trials), and **C** in human attention task (average across 100 attend out trials).

Given that beta is event-like in unaveraged data, there are several possible features of such events that could contribute to differences in prestimulus beta power averaged across time and frequency. The features that could underlie a low vs high value of averaged prestimus power (Figure 3 top) include a difference in: The number of prestimulus events (i.e. rate); event power; event duration; and/or their frequency span (Figure 3A-D, respectively). We analyzed the contribution of each of these features to the observed impact of trial mean prestimlus beta power on behavior as shown in Figure 1.

**Figure 3.**
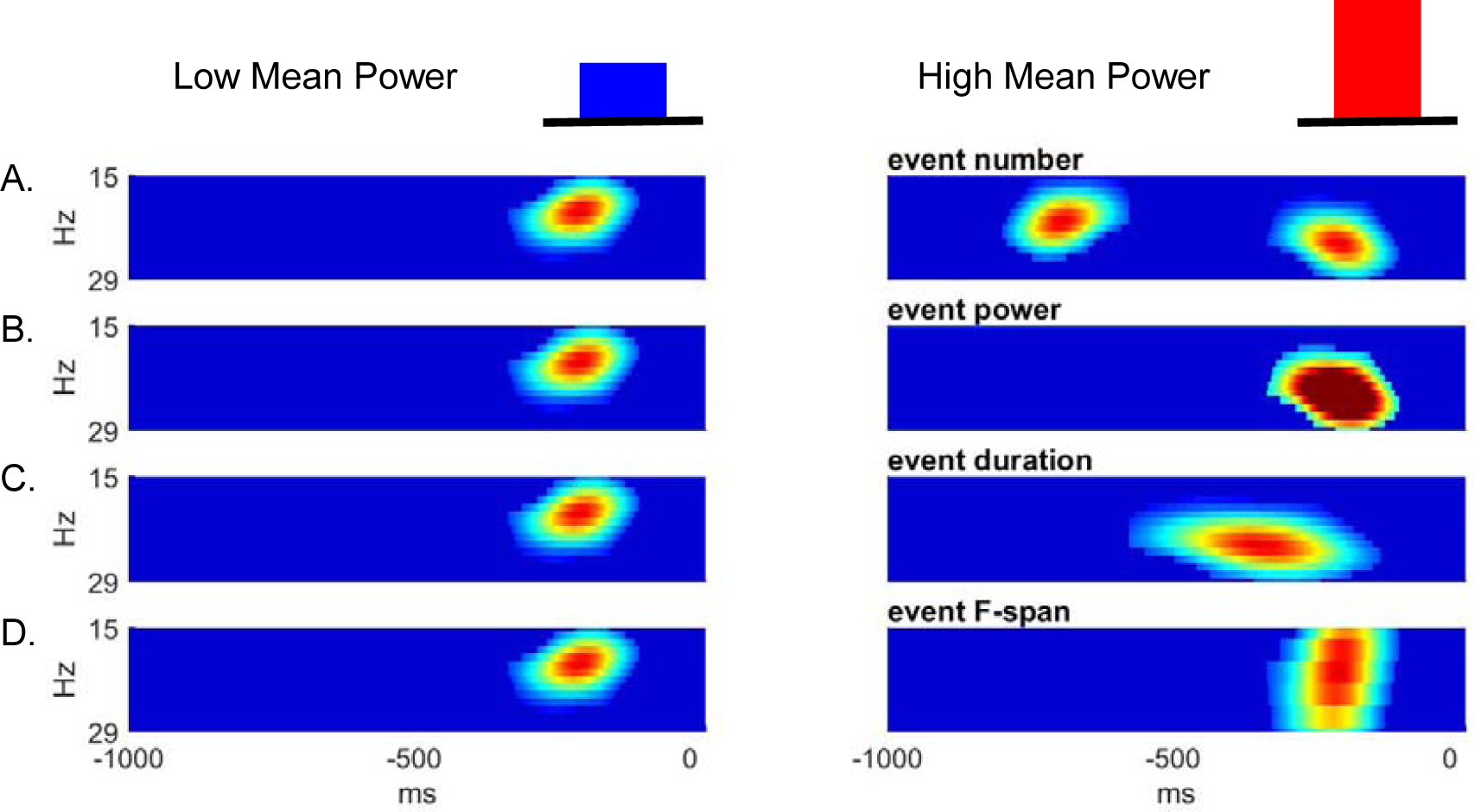
Schematic illustration of possible features underlying differences in averaged prestimulus beta power. Given that surges in beta power in non-averaged data occur as transient events, higher trial mean beta power could be due to an increase in A. event number (i.e. rate), B. event power, C. event duration, and / or D. event frequency span (F-span).

To quantify these high-power transient beta events in the non-averaged spectral activity, we defined beta events as local maxima in the 1 second prestimulus spectrogram for which the maxima frequency fell within the beta band (15-29Hz) and the maxima power exceeded a set power threshold. To choose the power threshold in a principled manner, we calculated for each subject the correlation between the percent area in 1 second prestimulus beta-band spectrogram above threshold (see red in Figure 4Ai inset) and the trial mean prestimulus beta power (Figure 4i). We chose the value of 6X median power as our threshold for further analyses because the correlation with mean power was highest near this point for most sessions across species and behavioral tasks (Figure 4i). The correlation peaking around 6X median power threshold suggested that the power above this threshold accounted best for the variability in trial mean prestimulus power; the focus of our study (see also Supplementary Figure 1 and Supplementary Text for examination of threshold variation). The inverse cumulative density function (1-CDF) of all local maxima in the beta band as a function of threshold shows that 6X median power threshold captures ∼20% of all local maxima (Figure 4ii).

**Figure 4.**
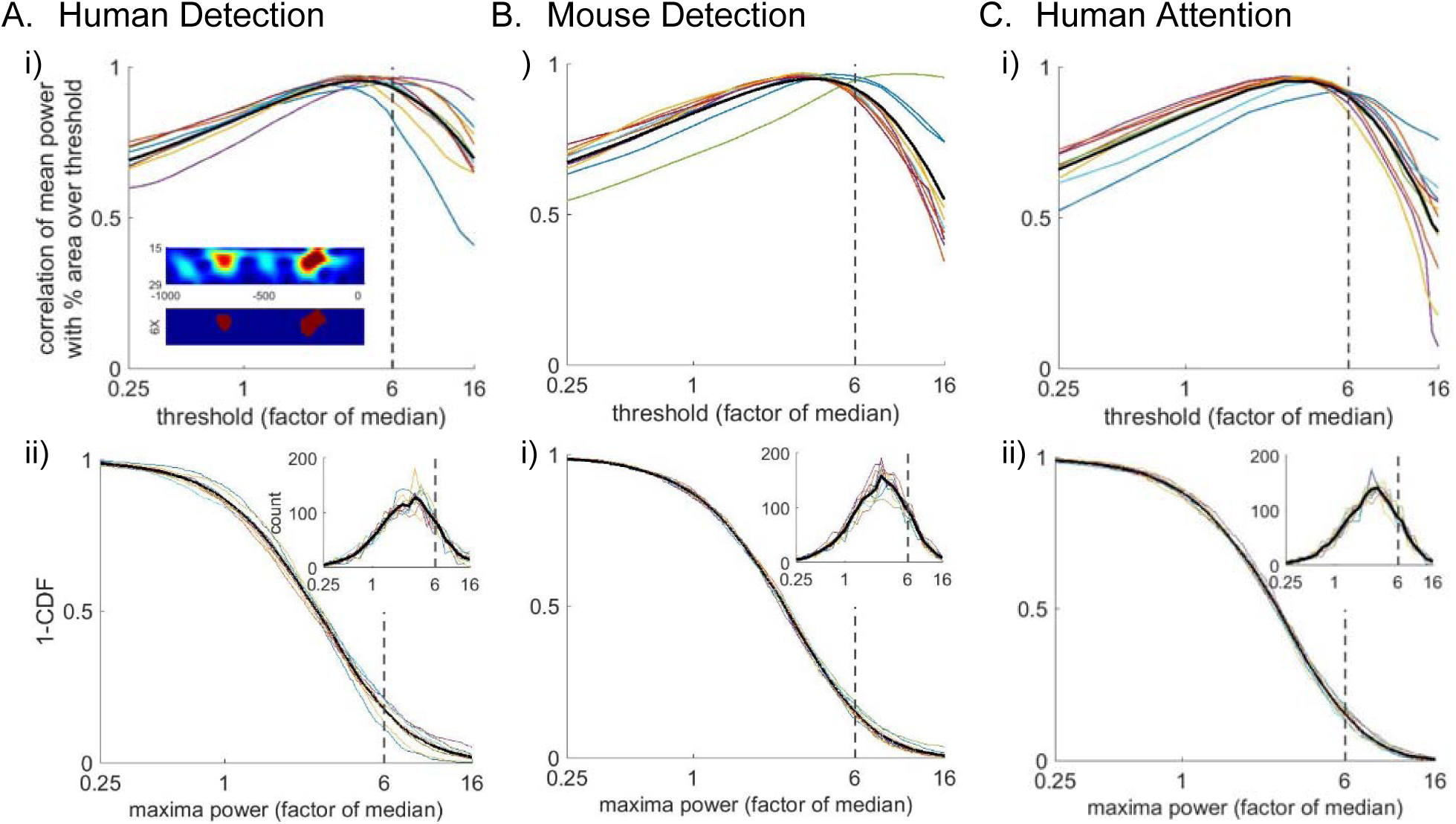
Beta events defined by a 6X median power threshold show consistently high correlation with average prestimulus beta. **i)** Trial-by trial correlation between mean prestimulus beta power and the percent area (i.e. percentage of pixels in the spectrogram) above threshold in the non-averaged spectrogram, for various thresholds calculated as factors of median and plotted on a log scale, in each data set (**A-C**). Inset in A shows an example spectrogram (top) with illustration of percent area above threshold for a 6X median threshold. **ii)** Distribution of maxima power for all beta-band local maximas in non-averaged spectrograms. 1-CDF shows proportion of local maxima that survived the thresholding (mean across trials for individual subjects / sessions are colored; grand mean across subjects is black). Insets show the same data as histograms.

Figure 5 displays the probability density function (PDF) of the number of events in the prestimulus period, and their mean power, duration and frequency span for each subject and task. Event power was defined as the spectrogram value at the maxima, and duration and frequency span were defined as full-width-at-half-maximum of the above threshold beta event in time and frequency dimensions, respectively. There was homology in the distribution of beta event features across tasks and species. Typically <4 beta events occurred in the 1-second prestimulus period; the event power PDF fell off monotonically after the power threshold; and event duration and frequency span were all confined to a restricted range around a stereotypical value (mean duration ∼110ms; mean frequency span ∼7Hz). Next, we assessed the impact of each of these features on mean prestimulus beta power, and on behavioral measures of perception and attention.

**Figure 5.**
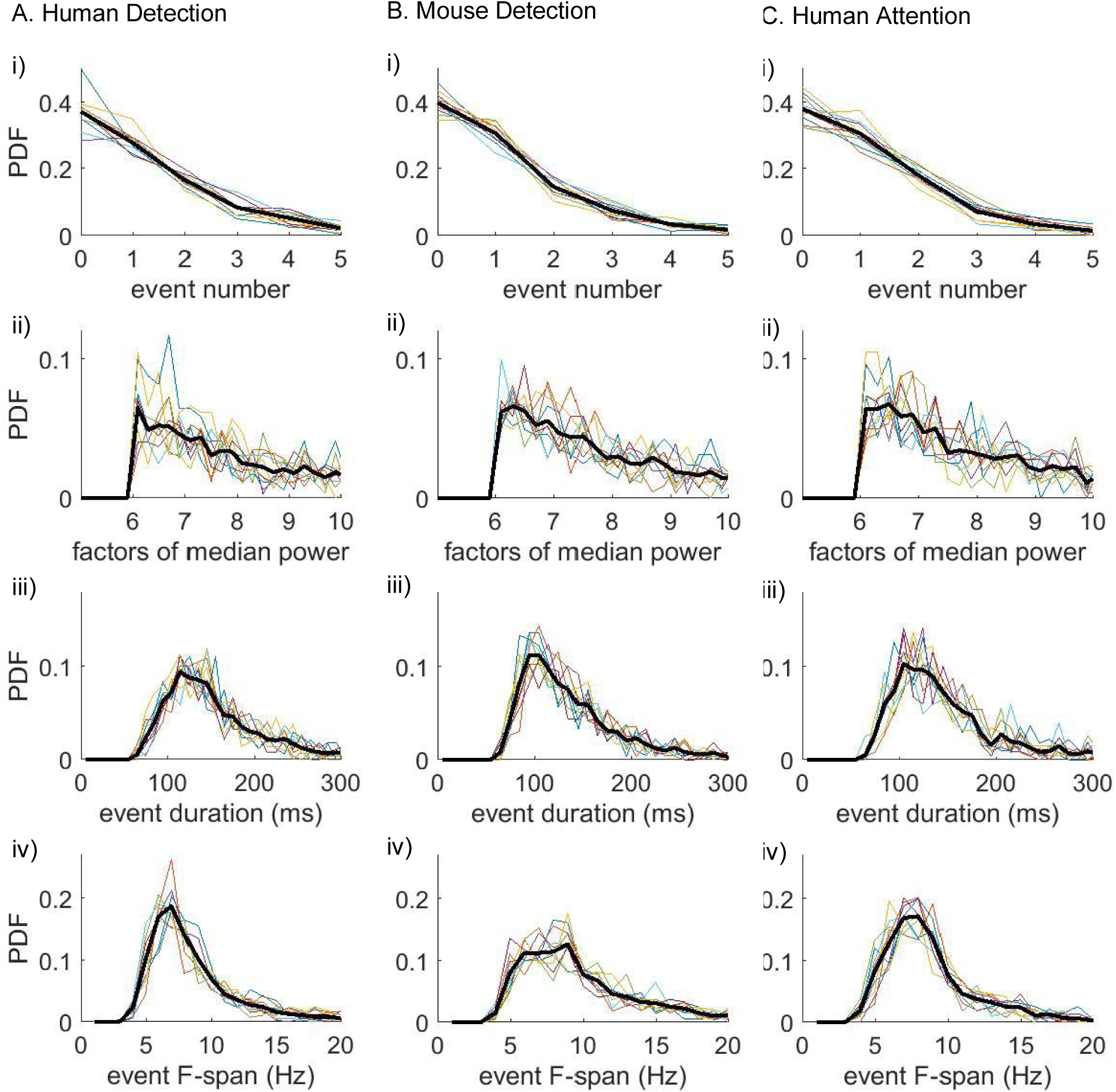
Beta event features are highly conserved across tasks and species. PDF for each prestimulus beta event feature; **i)** event number, **ii)** event power, **iii)** event duration, **iv)** event frequency span; aggregated across all trials, for each subject/session (colored) and the average across subjects (black), in each data set **(A-C)** using a 6X median threshold. Bin intervals for each PDF was as follows; 1 for event number; 0.2 (factor of median) for event power; 10 (ms) for duration; and 1 (Hz) for frequency span.

### The rate of prestimulus beta events had the highest correlation with trial mean prestimulus power and was the most consistent predictor of behavior

Figure 6 shows for all subjects and sessions in each dataset, the trial-by-trial correlation between trial mean prestimulus power and the trial summary of each beta event feature. Again, there was consistency across species and tasks. In each data set, the number of beta events in the prestimulus period (i.e. rate) showed a significantly higher correlation with averaged beta power than the trial mean prestimulus event power, duration or frequency span (asterisk Figure 6, see also Supplementary Figure 2 for regression analysis).

**Figure 6.**
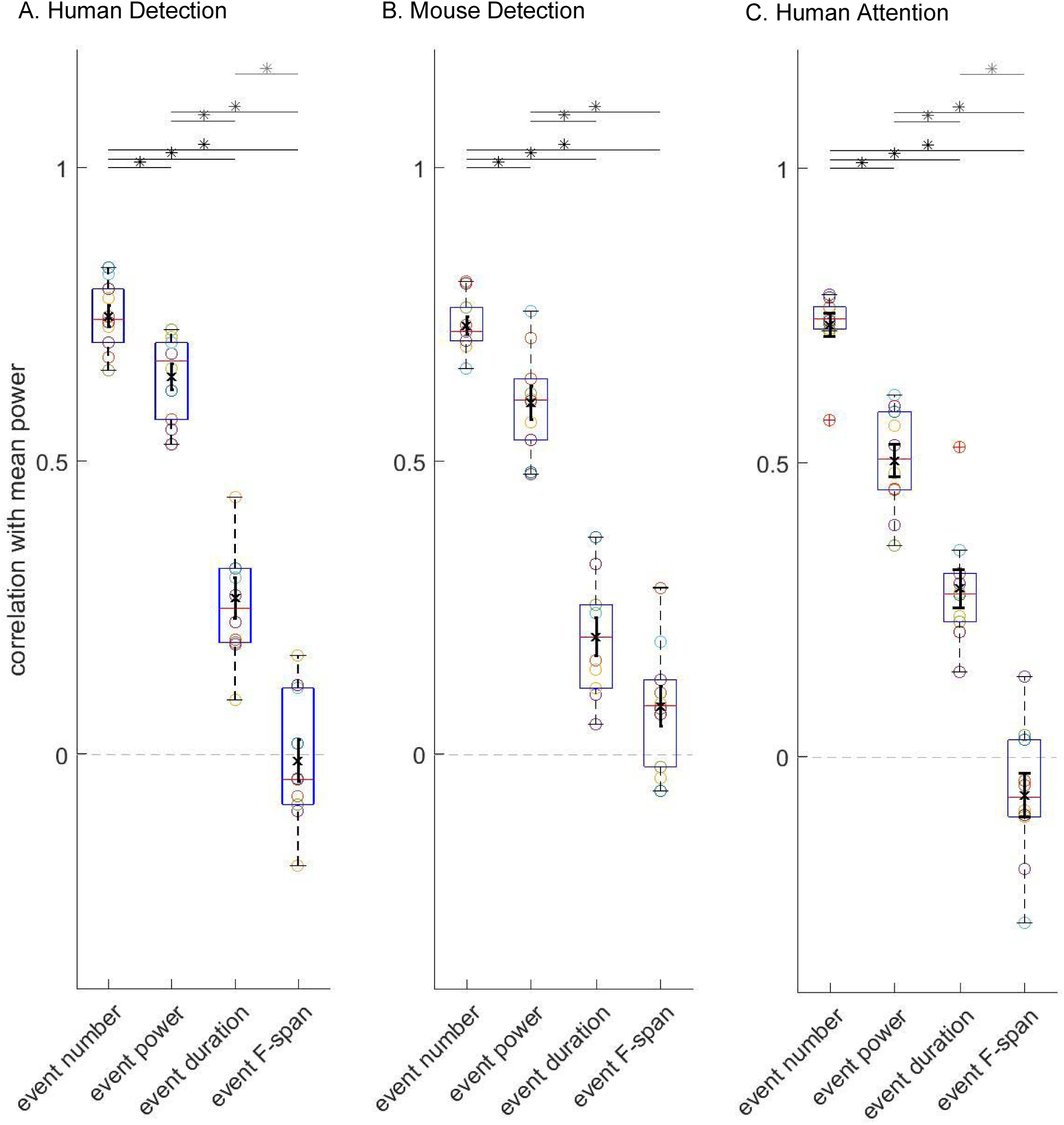
The number of beta events has a higher correlation with averaged prestimulus beta power than event power, duration or frequency span. Box and whisker plots over subjects / sessions depicting the Pearson's correlation coefficients (*R*) between each beta event features and prestimulus mean beta power, calculated on a trial-by trial-basis. Note, for all analyses involving trial mean event power, duration and frequency span, only the trials with one or more events were considered. The mean and ±SEM across subjects are plotted with a black x and error bars. Wilcoxon signed-rank test was performed for each pair of features and significance was determined at p<0.05 (asterisks). See also Supplementary Figure 2 for regression data.

The number of beta events in the prestimulus period was also the most consistent predictor of behavior across datasets. As in Figure 1i, we sorted the data from low to high values for each beta event feature (number, power, duration, frequency span) and calculated the corresponding percentage of detected or attend-in trials in PCM (Figure 7i). Several of the features showed a significant negative regression between feature value and detection or attention, such that the greater the value, the less likely the subject detected or attended to the stimulus (p<0.05). However, the *R*^2^values for event number were consistently higher across data sets.

**Figure 7.**
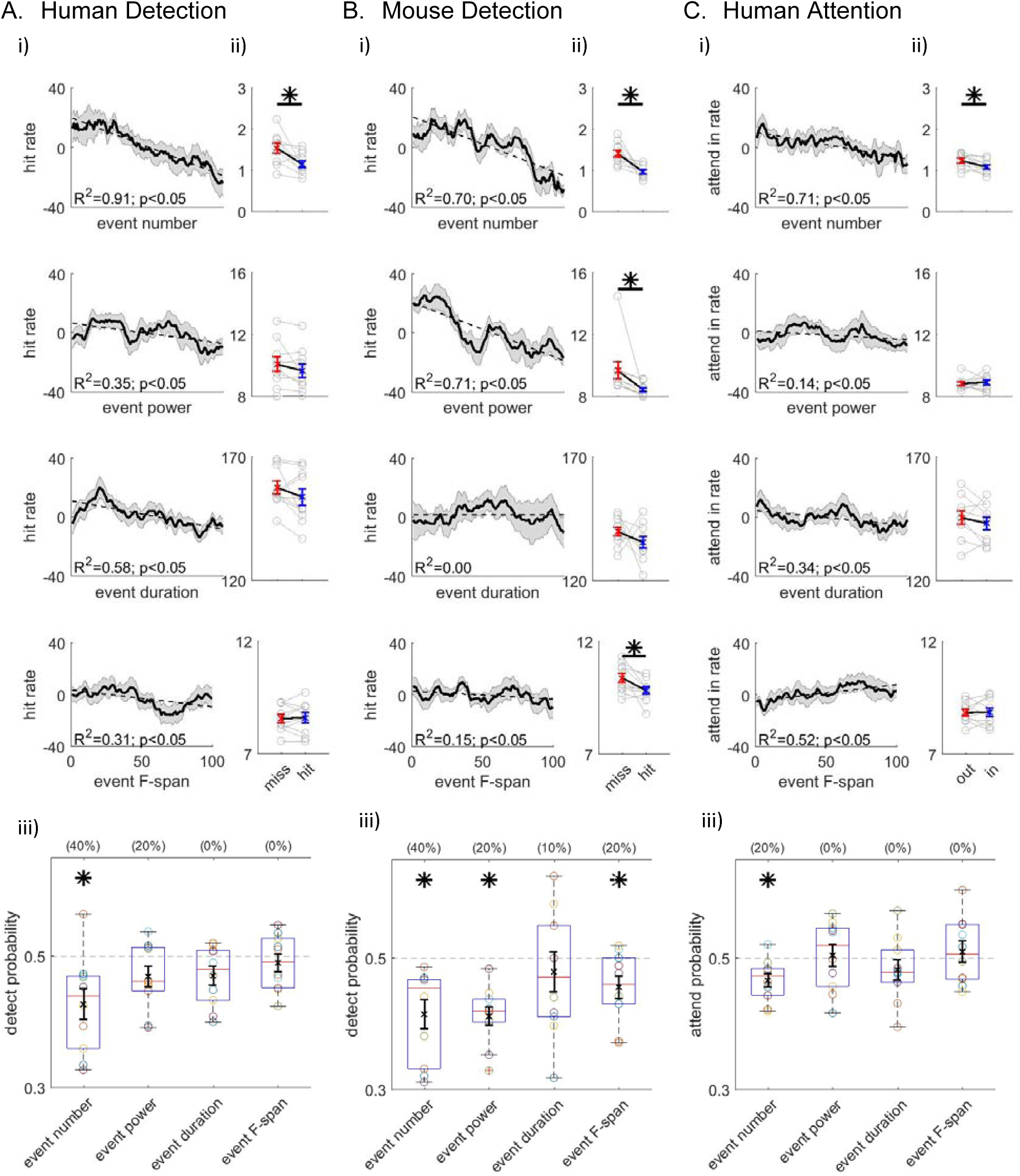
Beta event number (i.e. rate) consistently predicts detection/attention across tasks and species. As in Figure 1, where each analysis was performed separately for each event features (number, power, duration, frequency span), as labeled, for each data set **A-C**. **i)** Linear regression; **ii)** averaged data pooled across-subject / session and trials for each behavioral condition; and **iii)** box and whisker plot representation of DP / AP values across subjects/sessions. In the human data, only the number of events was predictive of detection / attention (DP / AP < 0.5). In the mouse data, event power and frequency span were also significantly predictive of detection, however the percentage of sessions showing significance was highest for event number (40% compared to 20%, see parentheses in **Biii**). Again, only the trials with one or more events were considered for trial mean event power, duration and frequency span.

Moreover, in the human data, when the grand average of each feature in the prestimulus period was pooled across conditions, only the number of events per trial was significantly higher in non-detected (miss) / attend-out conditions (Figure 7ii, asterisks p<0.05, two sample, left-tailed Wilcoxon signed rank test), and only the number of events was predictive of behavior when applying DP or AP analysis across subjects (Figure 7iii, asterisks p<0.05, one sample, left-tailed Wilcoxon signed rank test). The number of beta events was also predictive of behavior in the mouse LFP. In the LFP, event power and frequency span were also predictive of behavior. However, a smaller number of sessions showed significant DP<0.5 for event power and frequency span than the number of events, suggesting the rate of events in the prestimulus period was the most consistent predictor of behavior (40% for event number, and 20% for event power and frequency span, see parenthesis Figure 7iii, as determined by 95% bootstrapped confidence interval). The frequency span difference resulted from a differential 1/f effect in the mouse LFP data that did not exist in the human MEG (Figure 2i).

As stated above, we chose a 6X median power threshold to define beta events because this value consistently showed the highest correlation between percent area (in spectrogram) above threshold and averaged prestimulus power (Figure 4). We investigated the impact of varying the threshold on the correlation between each beta event feature and trial mean prestimulus power (Supplementary Figure 1i, see also Supplementary Text). We found that as the beta event threshold was lowered, event power became more strongly correlated with mean prestimulus power and event number less correlated (at ∼3X median threshold), as might be expected from including a greater number of peaks in the spectrogram that are within the beta frequency but smaller in amplitude and closer to the signal to noise boundary. Similarly, event power was a stronger predictor of behavior than event number for low thresholds (∼<3X median, Supplementary Figure 1ii, see also Supplementary Text).

### Non-detected trials were more likely to have a prestimulus beta event closer to the time of the stimulus

Given that the rate of prestimulus events influenced perception, we further hypothesized that the timing of the beta event within the prestimulus period could also impact perception. Calculating the PDF of the prestimulus event timing occurring closest to stimulus (i.e., most recent event) showed that non-detected trials had more events occurring close to the time of the stimulus than non-detected trials (Figure 8A/Bi). The PDF profiles were significantly different out to ∼300ms prestimulus in the human and mouse data, suggesting a dominant influences of beta events on function in this timescale (pointwise left-tailed Wilcoxon signed rank test, p<0.05). Note that the fall-off in PDF near the stimulus onset is due to edge effects of the spectral analysis when using Morlet wavelet convolution. The attention data did not have enough non-detected trials for analogous analysis (1).

**Figure 8.**
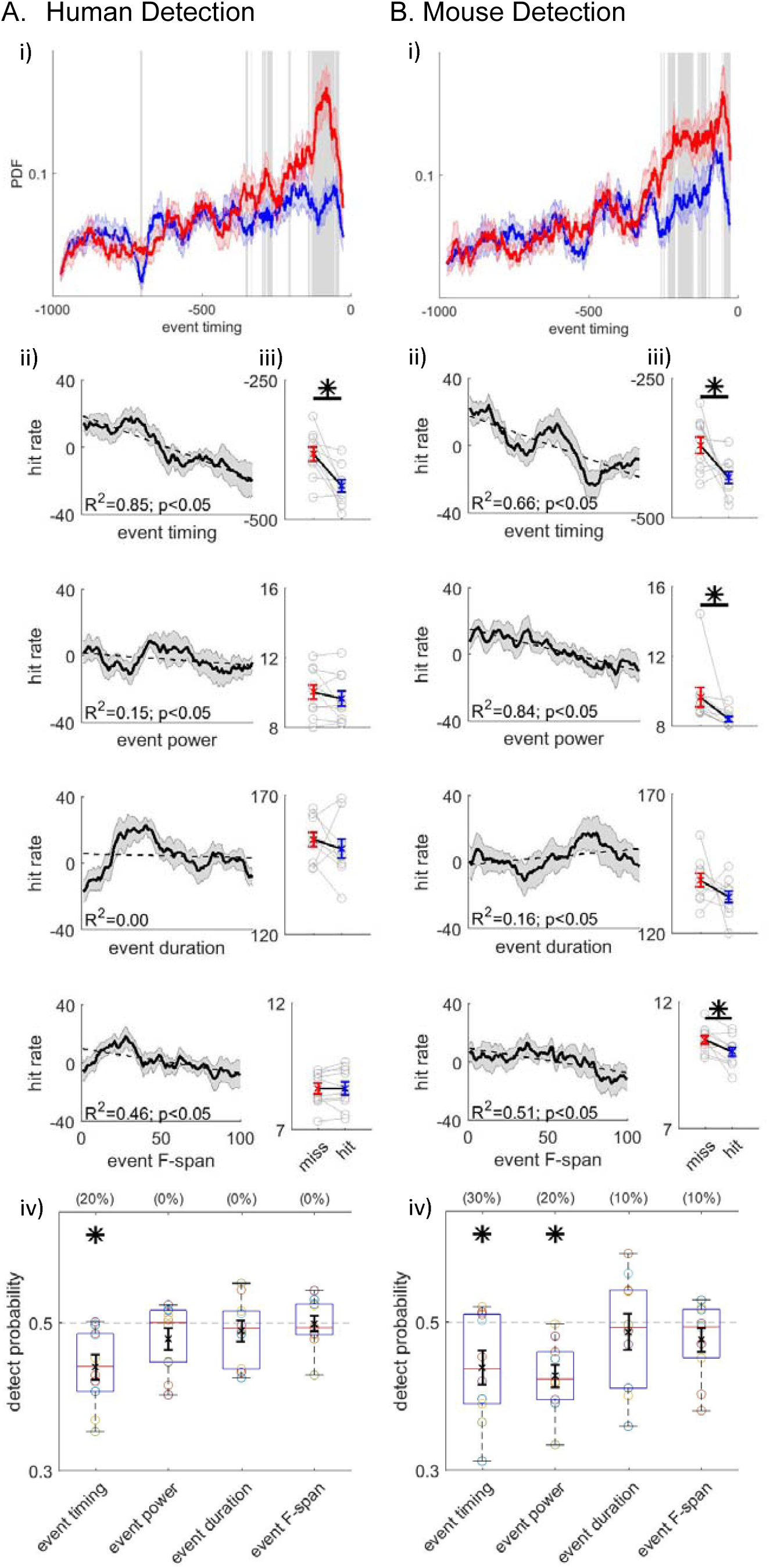
Non-detected trials are more likely to have a beta event closer to the time of the stimulus. **i)** PDF of the “most recent beta event” timing in detected (blue) and non-detected trials, in the human and mouse data (**A-B**) (mean ±SEM as error bars using 50ms windows sliding in 1ms steps. Grey shaded regions show significant differences at p<0.05 (pointwise left-tailed Wilcoxon signed rank test). Non-detected trials are more likely to have “most recent beta events” out to ∼300ms before the stimulus than detected trials. **ii-iv)** As in **Figures 1** and **7**, where each analysis was performed on features of the “most recent beta event” (event timing relative to the stimulus, power, duration and frequency span), as labeled. Only the timing of the most recent event relative to the stimulus was significantly predictive of detection in both data sets, and in the mouse data event timing showed a higher percentage of sessions significance DP<0.5 than event power on a per session basis (30% compared to 20%, see parentheses in **Biv**).

We further assessed the relationship between detection performance and several features of the most recent beta event, including its time relative to stimulus onset, and its power, duration and frequency span. Detection performance was quantified with the same three measures as in our prior analysis shown in Figure 7. In the human data, the most recent event timing relative to the stimulus onset was the only significant feature (left-tailed Wilcoxon signed-rank test of the average across behavior outcomes, and one-sample left-tailed Wilcoxon signed rank test of the DP distribution), such that the closer the most recent beta event was to the stimulus the less likely the subject was to detect (Figure 8Aii-iv). In the mouse data, the timing of the most recent beta event also predicted detection (Figure 8Bii-iv). Here, event power and frequency span also showed significant differences. However a smaller percentage of sessions showed significant DP<0.5 for event power and frequency span than for event timing, suggesting the timing of the most recent beta event was the most consistent predictor (30% for the most recent event timing, and 10% for event power and frequency span, see parentheses Figure 8iv). Notably, the choice of threshold for most recent event timing showed a similar trend to that of event number. As the threshold was lowered below ∼3X median, the most recent event power had a more significant influence on perception than the timing relative to the stimulus (Supplementary Figure 1iii, see also Supplementary Text).

### Beta events did not occur rhythmically

We next assessed whether beta events occurred rhythmically in the prestimulus period (i.e, with a regular Inter-Event-Interval, IEI), and if the degree of rhythmicity impacted behavior. The IEI distributions did not differ between detected or attend-in and non-detected or attend-out conditions (Figure 9i). Moreover, there was no clear evidence in the IEI distribution that beta events occur rhythmically in 1 second prestimulus windows. The lack of rhythmicity was further supported by calculation of the Fano Factor and the coefficient of variation (CV) of the beta event timings. For a Poisson process, the Fano factor and CV would be 1, whereas for a more regular (rhythmic) process, both values would be less than 1. In all three data sets, both values were around one or above (Figure 9ii). To further investigate the characteristics underlying the beta event generation process, we looked at the relationship between the Fano factor and CV^2^. A renewal process, defined as a process by which the time of occurrence of an event only depends on the time of occurrence of the previous event, would have similar values for Fano factor and CV^2^. However, in each data set, almost all data points had Fano Factors smaller than CV^2^, indicating that the driver underlying beta event generation was not a renewal process.

**Figure 9.**
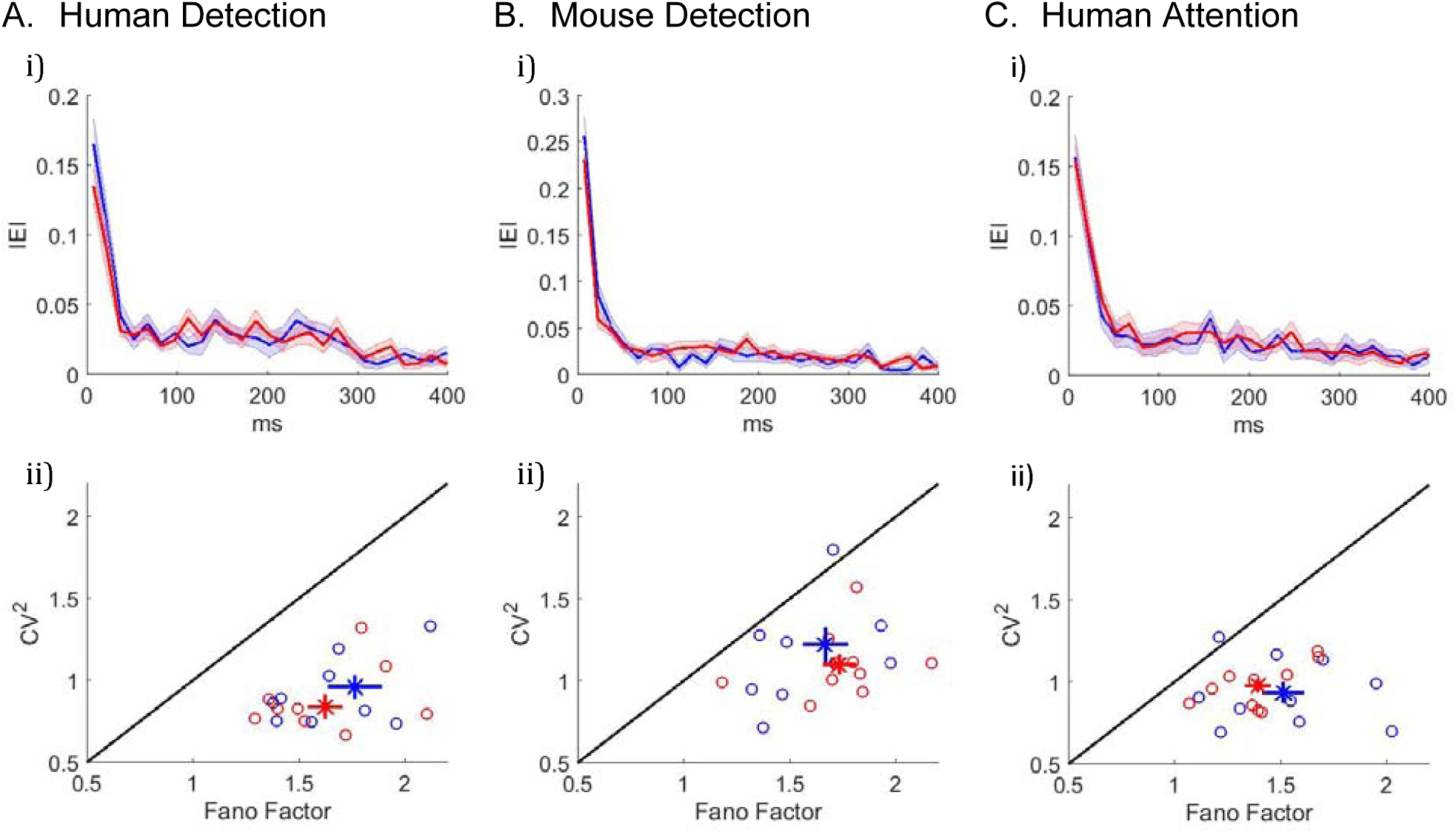
Inter-event-interval (lEI) analyses suggests that beta events are not rhythmic or a renewal process. **A** Human MEG SI detection data **B** Mouse LFP SI detection data **C** Human MEG SI attention data. **i)** IEI distribution (15ms bin intervals) on detected / attend-in (blue) and non-detected / attend-out (red) conditions; mean across subjects with ±SEM as error bars. **ii)** Fano factor and CV^2^ for each subject are plotted as ∘, for just the detected / attend-in trials (blue); non-detected / attend-out trials (red); and all trials (black). The mean across subjects are plotted as x, with and ±SEM for error bars.

## Discussion

Averaged prestimulus beta activity in somatosensory neocortex is predictive of perception and shifts in attention, such that higher averaged prestimulus beta power reflects decreased detection or attention. This result held across various analysis methods, species and recording modalities (human MEG, mouse LFP), consistent with the general predictions that spontaneous sensorimotor beta acts as, or is the by-product of, an “inhibitory” process (1, 3, 6, 28). To further uncover the potential mechanistic impact of beta on perception, we studied the trial-by-trial relationship between beta and behavior. On individual trials, beta emerged as a brief high-power transient above background noise in the frequency spectrum. As such, we postulated that observed differences in averaged power across behavioral conditions could reflect modulation in the number of high-power beta events in a predefined window (i.e. rate) and/or modulation in the event power, duration, or frequency span.

Our results showed that the rate of prestimulus beta events was the most consistent predictor of detection and shifts in attention, and most strongly correlated with average prestimulus beta power. In the human data, only the rate of events was significantly predictive of function across detection and attention tasks. Moreover, non-detected trials were more likely to reveal an event within ∼300ms of stimulus onset than detected trials. The timing of the most recent event was significantly predictive of perception, while the most recent event power, duration and frequency span were not. We did not find evidence that beta events occur rhythmically. The strong consistency across species and recording modalities suggests that differences in rate and timing of brief high-power prestimulus beta events is a general feature that underlies functionally relevant differences in averaged beta power.

Taken together, these results indicate that with shifts in attention, the brain decreases the probability of high-power beta events, and this decrease translates to improved sensory processing. This conclusion has direct implications for potential circuit level mechanisms subserving behavior. We conjecture below that the modulatory impact of beta events depends on their rate, such that the higher the rate of events near the stimulus, the greater the impact on perception.

### Relationship to prior studies

The finding that beta rhythms in neocortex are transient is consistent with many recent studies showing brief bouts of spontaneous beta band lasting <150ms in unaveraged data (18-21). In our own prior work studying human MEG, we have observed that the SI “mu” complex contains alpha and beta components that are brief in time and non-overlapping (17, 18, 22). Quantification of the highest power SI beta components showed that they typically lasted <150ms and had a stereotypical waveform. Moreover, beta activity from right inferior frontal neocrotex exhibited a similar profile (18).

Recent studies analyzing LFP in awake behaving non-human primates have shown that differences in averaged beta activity across behavioral conditions can reflect a difference in the accumulated density of transient beta “bursts” across trials (19, 20). Feingold and colleagues investigated movement induced changes in transient beta (13-30Hz) activity, termed beta “bursts”, in motor cortex and striatum from awake behaving monkeys (19). They found burst probabilities peaked at different times in different regions, reflecting differences in motor and cognitive demands. The LFP recordings in monkey putamen also showed bursts of beta under different motor demands, namely a synchronized-continuation task and reaction time task, such that the temporal alignment of bursts were task dependent (24). More recently, Lundqvist and colleagues showed that working memory is associated with transient bursts of 20-35Hz beta in prefrontal cortex (20). Beta burst characteristics in LFP have also been shown to be modulated by deep brain stimulation in the subthalamic nucleus of Parkinson’s disease patients (21).

### Homology in MEG vs LFP and implications for mechanism underlying beta event generation

Our data showed strong homology in beta event features in SI from human MEG and mouse LFP. We have also recently shown that, in both MEG and LFP, the highest-power beta events (maximum power events taken from 50 trials) had a stereotypical waveform shape where the frequency was determined by a dominant peak whose duration is one beta period (i.e. ∼50ms) (18). This homology may seem unexpected given the vastly different spatial scales over which MEG and LFP sample (29). Macroscale MEG oscillations are presumed to represent the summed sub-threshold activity across larger populations of neocortical pyramidal neurons than LFP. Consistency in beta event features across scales indicates that the high-power beta events represent the sub-threshold activity of large populations of pyramidal neurons picked up by both recording modalities.

In our prior modeling, converging evidence supported the theory that the circuit-mechanisms creating the highest power beta events consisted of bursts of sub-threshold excitatory synaptic drive targeting proximal and distal dendrites of pyramidal neurons, such that the distal drive was effectively stronger and lasted ∼50ms (18). Smaller populations of spiking neurons within beta events are likely present and influence behavior, but are subservient in the recordings to the larger scale events. Overall, the homology supports the analysis of high-power beta activity in translational research across animal models.

We defined beta events by choosing a threshold at which the power above threshold best accounted for the averaged beta power across the entire 1-second prestimulus window (6X median, Figure 4A) because our primary focus was to determine beta event features accounting best for functionally relevant difference is averaged prestmulus power. Our sliding threshold analysis showed that as the threshold was decreased and more low power beta maximum were considered, event power became a more consistent predictor of averaged prestimulus power and behavior than event number (near ∼3X median, Supplementary Figure 1). Based on the following, we conjecture that high power beta-events are generated by a distinct circuit-mechanisms that influences behavior.

For the 6X median threshold considered, beta events where within the top 20% of the highest-power beta events across all maxima in the beta band (Figure 4B), i.e., among those that have a stereotyped waveform shape (18). Further analysis of beta event features with different thresholds showed that as the threshold increased (>6X median), the duration and frequency span of the beta events become stereotyped, whereas for lower thresholds the distribution was far more widespread (see Supplementary Figure 3). These results suggest that, for larger thresholds defining high-power events, beta events represent a stereotyped process, whereas for lower thresholds they are more contaminated by “noise” that has high variability. Taken together, our prior and current results suggest that the mechanisms of the high power beta events that dominate fluctuations in the averaged prestimulus beta power are distinct in their generation and relationship to function.

An inter-event-interval analysis showed that beta events do not occur rhythmically. In fact, Fano Factors exceeding 1 in all subjects implies that the underlying process was less regular than a Poisson process, and more akin to a “bursty” process. This suggestion is in good agreement with our previous modeling studies predicting beta events emerging within the SI alpha/beta complex rhythm can be generated by repeated bouts of brief, burst-like excitatory synaptic drive targeting proximal and distal dendrites of pyramidal neurons (17, 18, 22). In these studies, the driving bursts were repeated approximately every 100ms, however stochasticity in their timing and effective strength created an alpha/beta complex rhythm, where beta emerged intermittently. Of note, all our analyses were restricted to 1 second prestimulus, which limited the analysis of rhythmicity, and rhythmic behaviors may be observed on longer timescales.

### Potential mechanisms underlying beta’s “inhibitory” impact on function: Rate vs timing

The precise role of brain rhythms in function is highly debated (30-34) (35-39). A well-established theory suggests that rhythms improve information processing via communication through coherence (40). This theory relies on the alignment of oscillatory phases between communicating areas, in which some phases are beneficial for the passing of information and others are not. In support, multiple studies have shown that during gamma oscillations created by reciprocal excitatory-inhibitory synaptic coupling there are “windows of opportunity” in which information can be transferred (e.g. (41, 42)). The phase of lower frequency alpha and beta oscillations have also been shown to correlate with spiking probabilities (43) and with perceptual benefits (44, 45).

Our data suggests that while phase alignment may be relevant in stimulus driven conditions where timed-locked signals (or responses) can align, it is unlikely that in spontaneous, prestimulus, anticipatory states beta’s local influence on the filtering of sensory input is through a temporal phase-alignment code. Rather, in all three of our tasks, the precise timing of the sensory stimulus onset is unpredictable for the subject (i.e. randomized design). Under these conditions, our results suggest that beta’s inhibitory correlation with perception may reflect the presence of another process that enhances beta event probability (i.e. rates), such as a period of specific neuromodulatory tone that promotes the drivers of this rhythm. Further, if a beta event occurs within 300ms before the sensory stimulus, the stimulus is unlikely to be perceived, suggesting that the increased rate in non-attended conditions acts to increase the probability that an event will occur close to, or perhaps during, the stimulus. The circuit and cellular mechanisms underlying beta event generation could in turn be the element undermining effective relay of information, as discussed below.

Coherence between spontaneous beta activity in distinct brain areas emerges in spontaneous states and predicts perception. As one example, we previously reported that prestimulus beta-band phase-locking values between SI and inferior frontal cortex (IFC) are different when comparing attended to non-attended states, such that attended trials exhibit smaller phase synchrony (10). Given our current results, smaller phase synchrony values may occur when the rate of beta events decreases in attended conditions.

### How might an increase in the rate of prestimulus beta events in SI translate mechanistically to a decrease in tactile perception?

One inference is that the circuit-level mechanisms creating beta events recruits an “inhibitory” process that decreases the relay of sensory information to or from SI. The more that beta events occur, the more likely this inhibitory process will be present at the time of the stimulus. In our recent work, converging evidence from modeling and human, mouse and monkey data showed that the stereotypical highest-power beta events were generated by “bursts” of excitatory synaptic drive targeting proximal and distal dendrites of pyramidal neurons, such that the distal drive was effectively stronger and lasted ∼50ms (18). Given this mechanism, the strong distal drive required for beta generation could also recruit inhibitory neurons in supragranular layers that act to decrease the efficacy of communication through supragranular channels (17). Alternatively, the generators of the exogenous drive contributing to beta emergence in SI (e.g. thalamic bursting) could mediate beta’s inhibitory impact on perception, in which case the increase in SI beta events represents an epiphenominal reflection of this external process. Future studies are necessary to test alternative hypotheses and to identify possible cortical or thalamic sources of the exogenous drive.

### Implications for future studies

An open question is if properties of transient rhythms in other frequency bands may underlie observed differences across behavioral conditions. Indeed, while many rhythms are sustained for several cycles (e.g. occipital alpha rhythms during eye closure, slow wave sleep rhythms), transient rhythms emerging for a few cycles have been reported in other bands, including gamma (20, 46, 47) and alpha (17, 18, 22). Lundqvist and colleagues reported that gamma burst rates increased with memory load in a working memory task, and firing in neurons predictive of encoding and correlated with changes in gamma burst rates (20). We predict similar results will hold for alpha events as for beta events.

The transient nature of high-power spontaneous beta events has strong implications on brain stimulation studies, such as transcranial alternating current or magnetic stimulation (tACS, TMS), aimed at entraining beta “rhythms” to causally modulate behavior. Causal manipulations may be more effective if they are designed to match the intermittency and temporal characteristics of the beta events observed in real behavior.

Overall, our study highlights that when rhythms are transient in nature, signal differences in several domains (e.g. number, amplitude, duration, frequency span) could be the source of correlation between average power and function. Consistency in beta event characteristics across tasks, recording modalities, and species, suggest modulation of beta event rate may be the fundamental feature underlying behaviorally relevant differences in averaged beta power in a wide range of studies.

## Acknowledgements

We thank Christopher Black for help with figures. This work was supported by the US National Institutes of Mental Health (R01MH106174; S.R.J.); the Department of Veterans Affairs, Veterans Health Administration, Office of Research and Development, Rehabilitation Research and Development Service, Project N9228-C (S.R.J.); US National Institute of Neurological Disorders and Stroke (R01NS045130; C.I.M.); and fellowships from Fulbright and the Brown Institute for Brain Sciences to H.S.

## Author Contributions

S.R.J. and C.I.M. designed and S.R.J. conducted the human detection and attention experiments. H.S. and C.I.M. designed the mouse detection experiment and H.S. conducted mouse surgery and behavior training. H.S., R.L. and S.T. analyzed the data. H.S. and S.R.J. made the figures and S.R.J. H.S. R.L and C.I.M. wrote the manuscript.

## Methods

### Human Recordings

#### MEG data collection

Details of the human MEG recordings and source localization for both the detection and attention tasks have been previously reported (25, 26). In brief, we used the 306-channel Vectorview system. For both the detection and attention dataset signals were sampled at 600 Hz with the band-pass set to 0.01 to 200 Hz. Dipole activity of the postcentral gyrus in the hand area of SI was extracted by using least-squares fit with a dipole forward solution calculated using individualized structural MRIs, or a spherically symmetric conductor model of the head (48).

#### Detection task

Subject recruitment, experimental protocol, and data acquisition have been described in prior reports from our group (25, 26). In brief, the stimulus was a single cycle of a 100 Hz sine wave (i.e. 10ms duration) generated by piezoelectric benders (Noliac). The stimulus was applied to the third digit fingertip of the right hand. Individual subjects’ perceptual thresholds were obtained before imaging by employing a parameter estimation by sequential testing (PEST) convergence procedure (49, 50), which estimated the threshold to <5μm precision.

During MEG imaging, 70% of the trials were maintained at perceptual threshold (50% detection) using a dynamic algorithm. 10% suprathreshold stimuli (100% detection) and 20% null stimuli (catch trials) were randomly interleaved with the threshold stimuli. Trial duration was 3000 ms. Trial onset was indicated by a 60 dB, 2 kHz auditory cue delivered to both ears for a duration of 2000 ms. The stimulus was delivered at a random time between 500 and 1500 ms after the onset of the auditory cue. The subjects were given 1000 ms to report detection or non-detection of the stimulus with button presses using the second and third digits of the left hand, respectively.

Ten subjects underwent 8 runs, 120 trials each, in one data collection session. To minimize within-session training effects, we limited our analysis to the last 100 trials of perceived and nonperceived stimuli for each subject (51).

#### Cued attention task

Subject recruitment, experimental protocol, and data acquisition have been explained in detail previously (25). In brief, 10 subjects were instructed to fixate on a cross on a projection screen. Each trial lasted 3500ms, and began with a 60dB, 2kHz tone delivered to both ears, simultaneously accompanied by the fixation cross changing into a visual word cue instructing the subject where to attend; either “Hand” (attend-in condition), “Foot” (attend-out condition) or “Either” location. The tactile stimulus was delivered to the cued location at a randomized time between 1100 ms and 2100 ms after the cue onset. The stimulus was a single cycle of a 100 Hz sine wave (10 ms duration) generated by piezoelectric benders, as in the detection task. Stimuli were applied to the distal pads of the third digit of the left hand or first digit of the left foot, and PEST procedure (49, 50) was employed before the task to estimate subject’s initial detection threshold for both the hand and the foot. Both the auditory tone and the visual cue ceased after 2500 ms. Subjects then had 1000 ms to report detection or non-detection of the stimulus at the cued location, with button presses using the second and third digits of the right hand, respectively.

The task consisted of at least 5 cued detection runs, where each run consisted of 40 of each attention condition randomly intermixed (i.e. 120 trials per run), in one data collection session. Similar to the detection task, all analyses were limited to the last 100 perceived trials of attend-in and attend-out conditions for each subject. Note, limiting to perceived trials allowed us to dissect attention effects independent of detection performance.

### Mouse Recordings

#### Chronic extracellular electrophysiology in mice

All electrophysiology data was collected using the Open Ephys system. Continuous data was sampled at 30 kHZ, and off-line downsampled to 1000 Hz. To reject common noise shared across channels, such as muscle artifact, independent component analysis (ICA; https://research.ics.aalto.fi/ica/fastica/code/dlcode.shtml) was applied to the downsampled continuous data collected across all 64 channels throughout the entire duration of the session. Components resembling artifact were manually chosen, and denoised signal was reconstructed with the exclusion of the rejected components. This reconstructed signal was used for all further analyses. As all electrodes were located in the barrel cortex and showed highly correlated LFP, one electrode was chosen for each mouse for all analyses.

#### Detection task

##### Animals

Two neurologically healthy male mice were used in this experiment (5 sessions from each mouse). Mice were 8-15 weeks at the time of surgery. Animals were individually housed with enrichment toys and maintained on a 12-hour reversed light-dark cycle. All experimental procedures and animal care protocols were approved by Brown University Institutional Animal Care and Use Committees and were in accordance with US National Institutes of Health guidelines.

##### Surgical procedure

Naive mice were induced with isofluorane gas anesthesia (0.5–2% in oxygen 1 L/min) and secured in a stereotaxic apparatus. We injected slow-release buprenorphine subcutaneously (0.1 mg/kg; as an analgesic) and dexamethasone intraperitoneally (IP, 4 mg/kg; to prevent tissue swelling). Hair was removed from scalp with hair-removal cream, followed by scalp cleansing with iodine solution and alcohol. Then, skull was exposed by incision. After the skull was cleaned, muscle resection was performed on the left side. A titanium headpost was affixed to the skull with adhesive luting cement (C&B Metabond). Two small stainless-steel watch screws were implanted in the skull plates; one anterior to bregma, one on the right hemisphere. Next, a ∼1.5 mm–diameter craniotomy was drilled over barrel cortex of the left hemisphere, and subsequently duratomy was performed. The guide tube array (8 by 2 arrangement of 33ga polyimide tubes; 2 mm by 0.5 mm) was centered at 1.25 mm posterior to bregma and 3.25 mm lateral to the midline and angled 45 degrees relative to midline. The drive body was angled 30 degrees relative to the perpendicular direction to compensate for the curvature of barrel cortex. Once the implant was stably positioned, C&B Metabond and dental acrylic (All for Dentist) was placed around its base to seal its place. A drop of surgical lubricant (Surgilube) prevented dental acrylic from contacting the cortical surface. Mice were given at least 3 days to recover before the start of water restriction.

##### Trial structure and behavior control

The behavioral task setup was adapted from a previous study from our group (41) with slight modifications. Each trial consisted of right-side (contralateral) vibrissae stimulation in the dorsoventral direction, with 20Hz deflections that lasted 500ms (10 pulses with the same amplitude). The stimulus was delivered through piezoelectric benders (Noliac). Vibrissae on the right side were tied up with a suture loop fed through a glass capillary tube (0.8 mm outer diameter) attached to the piezoelectric bender. Most of the vibrissae were secured about 3 mm from the mystacial pad.

If the mouse licked within 700ms relative to the onset of the stimulus, a drop of water was delivered, shortly followed by vacuum suction to remove any remaining water not consumed by the animal. Water delivery and vacuum suction was controlled by a solenoid valve (NResearch) connected with Tygon tubing. Mice received water through a plastic tube, which was positioned near the animal’s mouth using a Noga arm. The water was delivered based on gravitational flow and the volume was controlled by the duration of valve opening (∼100 ms), calibrated to give an ∼3 μl per opening. Individual licks were detected using beam breaks of IR detector flanking the lick tube.

All behavioral events including piezoelectric control, reward delivery, and lick measurements were monitored and controlled by custom software written in Matlab and interfaced with a combination of Arduino and PCI DIO board (National Instruments).

##### Behavioral training

Mice were water restricted for at least 7 days before start of training, during which time mice were acclimated to the head-fixed setup where mice could freely run on a fixed-axis styrofoam ball. Mice were given at least 1ml per day, calibrated such that mice would not lose weight further than 80% of their original weight before water restriction.

Training began with reward-all session where vibrissae stimuli were paired with water delivery regardless of the animal’s response. This allowed mice to establish an association between vibrissae deflection and reward, such that the mice learned to lick when they detected the stimulus in anticipation of the reward. Mice learned the association after about a week of reward-all training. After this period reward was only delivered on trials where the mouse licked within 700ms of the stimulus onset (i.e. reward window).

The stimulus amplitude was varied on a trial by trial basis between 0 to maximal amplitude (about 1mm deflection) in a randomized manner. Before the start of each session, the experimenter set the percentage of maximal trials and 0 amplitude (catch) trials. The rest of the trials were submaximal trials, where the stimulus amplitude was randomly drawn from a uniform distribution between 0 and maximal amplitude. Throughout training, percentage of maximal amplitude trials were gradually lowered to 10%, and percentage of catch trials were gradually increased to 25%.

Licking during catch trials (false positives) lead to a time out of 15 seconds. Further, exploratory licking during inter-trial intervals (ITI, 2-6 seconds) lead to resetting (prolongation) of ITI, and this reset could happen up to 10 times. This prevented excessive impulsive licking.

If mouse did not consume enough water during the behavior session, supplementary water was given several hours after the conclusion of the session, such that mouse would have drank at least 1ml of water each day. Mouse weight was monitored throughout the entire duration of training such that their weight would not fall below 80% of their original post-surgery weight.

##### Behavioral analysis

Threshold trials were chosen offline from periods within the ∼2.5 hour session where the mouse was engaged in the task. Even in well trained mice, there were periods where animals defaulted to non-optimal strategies, such as excessive impulsive licking or non-engagement from satiety or prolonged inattention. To address the issue of exploratory licking that was not in response to detection, hit trials were defined as trials where the animal not only licked within the 700 ms reward window but also did not lick the spout prestimulus up to -1000 ms. In addition, we filtered out the trials between two false positives if there were no misses or correct rejections in between, to exclude periods where the mouse was acting impulsively. To filter out periods where the mouse was not engaged in the task, we looked at trials with strong stimuli (as defined as stimulus at 80% detection, calculated from Boltzmann distribution fit of psychometric performance over the entire session). If three consecutive strong stimuli lead to non-detection and there were no detected trials in between, all trials in between were filtered out.

We chose amplitude-matched threshold level detected and non-detected trials for all analyses. To do this, we binned all submaximal stimulus amplitude into 15 bins. Detected and non-detected trials were chosen from the high-performance periods to maximize trial count while matching the stimulus amplitude histogram.

### Common Data Analysis Procedures for Human and Animal Recordings

#### Spectral analysis and trial mean prestimulus beta power calculation

For all three tasks, we analyzed for each trial the -1 to 0 second window relative to the stimulus onset. The spectrograms of the spontaneous data were calculated for each prestimulus window by convolving the signals with a complex Morlet wavelet of the form 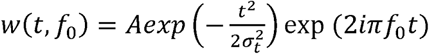, for each frequency of interest *f*_0_, where σ = *m*/2 π*f*_0_ and *i* is the imaginary unit. For both the human datasets, the spectrogram was calculated from 1 to 60 Hz, whereas for the mouse dataset, the spectrogram was calculated from 1 to100 Hz. The normalization factor was 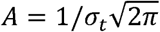. Consistent with prior publications from our group (18), we chose 7 for constant *m* (number of Morlet wavelet cycles). Time-frequency representations of power (TFR) were calculated as the squared magnitude of the complex wavelet-convolved data.

To compare TFR values across subjects / sessions, TFR was normalized by the median power value for each frequency. This median was calculated from all power values, at each frequency, in the -1000 to 0 ms prestimulus TFR concatenated across trials. Normalized TFR values are calculated in factors of median for each frequency, separately for each subject / session. For all analyses, the normalized TFR was used, such that the unit of all TFR values was in factors of median.

For trial mean prestimulus power, the normalized prestimulus TFR was averaged across time (-1000 to 0 ms window) and frequency band (15 to 29 Hz, inclusive), in each trial.

### Defining Beta Events and Features

#### Thresholding to define beta events

Beta events are defined as local maxima in the trial-by-trial TFR matrix for which the frequency value at the maxima fell within the beta band (15-29 Hz) and the power exceeded a set threshold.

To choose he power threshold that best captures variability in trial mean prestimulus power, we calculated the percent area in 1 second prestimulus beta-band spectrogram above threshold. That is, for the 1 second prestimulus beta-band TFR matrix in each trial, we quantified the percentage of matrix elements (i.e. pixels in spectrogram) that has power above threshold. For all analyses shown, except Supplementary Figures 1 and 4, the power threshold was set to be 6X the median power.

#### Beta event number

Beta event number was calculated as the number of beta events in the -1000 to 0 ms window for each trial.

#### Beta event power

The trial mean event power was defined as the power of all events averaged in the -1000 to 0 ms prestimulus period. For all analyses of trial mean event power, only trials that had at least one event were considered.

#### Beta event duration and frequency span

Beta event duration and frequency span were defined as full-width-at-half-maximum from the beta event maximima in the time and frequency domain, respectively. Edge cases in the time domain were handled in the following way: for maxima happening near the edge time points (-1000 ms and 0ms), if the power did not fall below the half-maximum at the edge, event duration was calculated by doubling the half-width of the side that was not cut by the relevant edge. The same method was used to handle edge cases for frequency span; if the power did not fall below half-maximum at either the lower boundary (1 Hz) or at the upper boundary (60 Hz for human, 100 Hz for mice), frequency span was calculated by doubling the half-width of the side that was not cut by the boundary.

The trial mean event duration and frequency span were defined as the averaged values in the 1-second prestimulus window. For all analyses of trial mean beta event duration and frequency span, only the trials that had at least one event was considered.

#### Most recent event timing and other features

The “most recent event” was defined as the event that happened closes to the stimulus onset in the -1000 to 0 ms prestimulus window. Most recent event timing was defined as the timing of the maxima of that event. Other features of the most recent event (power, duration and frequency span) were calculated as defined above. For all analyses involving the most recent event, only the trials that had at least one event was considered.

The PDF of most recent event timing in figure 8i) was calculated in 50ms windows sliding in 1ms steps. For each window, a left-tailed Wilcoxon signed rank test was applied across subjects / sessions to test whether the most recent event was more likely to happen within that window on non-detected trials. Note that the fall-off near the onset of the stimulus is due to edge effects in the TFR: Because of the tapering of power in wavelet-convolved spectrograms, local maxima near the temporal edges were less likely to be identified as high-power events. Therefore, we do not expect these fall-offs to be biologically grounded.

#### Regression Analysis

For each linear regression analysis, we sorted all trials in increasing order of the variable under consideration (e.g. prestimulus trial mean power). We then calculated the detection / attention rate in boxcar windows of 21 trials, slided in steps of 1; where the first bin was 21 bottom-ranking trials and the last bin was 21 top-ranking trials. If multiple trials had the same value (e.g. multiple trials with 1 event), the sorted trial order was shuffled to prevent artificial correlation. Detection / attention rate was then normalized as percent change from mean (PCM).

The human data was analyzed across 100 trials per condition in a single session, as described above. The mouse data had an uneven number of trials across sessions. Therefore, we used the Matlab “interp1” function to resample 100 bins of the detection / attention rate. The resampled detection / attention rate were averaged across sessions, then regressed across bins (calculated in percentiles of 100) using the Matlab function “fitlm”. The same analysis was applied to the human data. Significance of the linear regression was determined with the F-test.

#### Grand Average Analysis

For each grand averaged analysis reported, all prestimulus data under consideration for a given behavioral outcome / condition were averaged. Only trials with at least one event were considered, with the exception of trial mean prestimulus beta power and event number. Left-tailed Wilcoxon signed-rank test was used to test whether the median across subject / sessions was significantly higher for pooled averages of non-detected / attend-out trials.

#### Detect Probability / Attend Probability Analysis

Detect probability (DP) / attend probability (AP) was calculated as the area under a receiver operating characteristic (ROC) curve, where the ROC curve was defined by applying a binary classification of each variable under consideration (e.g. trial mean prestimulus beta power) to a behavioral outcome / condition (i.e., detection / attention). The area under the ROC curve analysis was performed with the Matlab function “perfcurve”. For all analyses, only trials with at least one event were considered, with the exception of trial mean prestimulus beta power and event number.

DP / AP of 0.5 indicates that the beta event feature under consideration cannot dissociate between the behavior outcome / condition. DP / AP under 0.5 signifies that beta event feature under consideration is significantly predictive of non-detected / attend-out trials. To quantify the percentage of significant subjects / sessions, a 95% confidence interval was determined individually for each subject / session by bootstrapping 10,000 times. If the upper boundary of the 95% confidence interval was below 0.5, the beta event feature under consideration was considered significantly predictive of non-detected / attend-out trials for that subject / session.

#### Inter-Event Interval (IEI)

The IEI was calculated as the time difference between two consecutive events, in trials that had two or more events. Note that because of the 1000ms window limit, IEI exceeding the window size could not be captured. The theoretical lower limit (resolution) of IEI was determined by the 7-cycle Morlet wavelet we used to generate the TFR matrices. All IEI values were pooled across trials within subjects / sessions, and the PDFs were calculated in 15ms bins.

#### Coefficient of Variation (CV^2^) and Fano Factor (FF)

Coefficient of variation of the inter-event intervals was defined as follows:

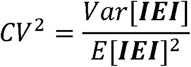

where *var*[*IEI*] is the variance, and *E*[*IEI*] is the mean.

Fano factor was used to quantify the trial-to-trial variability of event number per trial (EpT), and was defined in the following way:

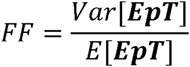

Renewal processes are defined as point processes with independent and identically distributed waiting times. Renewal processes are characterized as having equal CV^2^ and FF values for appropriately long trial lengths. Our results are limited by the fact that all analysis is restricted to -1000 to 0 ms prestimulus.

